# Identification of long non-coding RNAs as substrates for cytoplasmic capping enzyme

**DOI:** 10.1101/2022.07.17.500354

**Authors:** Avik Mukherjee, Safirul Islam, Rachel E. Kieser, Daniel L. Kiss, Chandrama Mukherjee

## Abstract

Cytoplasmic capping returns a cap to specific mRNAs thus protecting uncapped RNAs from decay. Prior to the identification of cytoplasmic capping, uncapped mRNAs were thought to be degraded. Here, we test whether long non-coding RNAs (lncRNAs) are cytoplasmic capping enzyme (cCE) substrates. The subcellular localization of fourteen lncRNAs associated with sarcomas were examined in U2OS osteosarcoma cells. We used 5’ rapid amplification of cDNA ends (RACE) to assay uncapped forms of these lncRNAs. Inhibiting cytoplasmic capping elevated uncapped forms of selected lncRNAs indicating a plausible role of cCE in targeting them. Analysis of published cap analysis of gene expression (CAGE) data shows increased prevalence of certain 5’-RACE cloned sequences suggesting that these uncapped lncRNAs are cytoplasmic capping targets.

## Introduction

The N^7^methyl guanosine (m^7^G) cap is added to RNA polymerase II (pol II) transcribed RNAs by RNA guanylyltransferase and 5’-phosphatase, commonly called mRNA Capping Enzyme (CE hereafter) bound to the phosphorylated C-terminal domain of pol II [1, 2]. CE’s N-terminal triphosphatase domain removes the terminal phosphate group from the first transcribed nucleotide of nascent transcripts generating an mRNA with a 5’-diphosphate end. CE’s C-terminal guanyl transferase domain then transfers GMP onto the diphosphate RNA end. Finally, RNA guanine-7 methyltransferase (RNMT), working as a heterodimer with RNMT-associated mini protein (RAMAC), methylates the N^7^ position of the newly-added guanosine to complete the m^7^G cap [3]. The m^7^G cap is recognized by a large number of proteins that regulate different stages of RNA metabolism including co-transcriptional processing, nuclear export, translation, and decay [4–7]. The m^7^G capped RNA can be further modified by methylating the 2’-hydroxyl of the RNA’s first (cap 1) and second (cap 2) transcribed nucleotides [8, 9].

Initially, decapping was thought to irreversibly consign an RNA to degradation. However, several studies have detected stable uncapped transcripts in different organisms including yeast, plants, parasites, and mammals [10–15]. The discovery of a cytoplasmic pool of CE (cCE) in mammalian cells offered a mechanism by which uncapped RNAs could be recapped [16]. Different forms of cCE are found exhibiting functions similar to mammalian cCE, as they have been demonstrated in Drosophila and Trypanosome [11, 17], suggesting that cCE is a regulator of post-transcriptional RNA control.

In contrast to nuclear CE, cCE functions in a complex of proteins where Nck1 acts as the scaffold to form the cytoplasmic capping complex [18]. RNAs can enter the cytoplasmic capping pipeline at two points. For RNAs with monophosphorylated ends, an unidentified 5’ RNA kinase adds a phosphate to the RNA generating a 5’-diphosphorylated RNA, which is the substrate of cCE [16, 18]. Alternatively, the substrate RNA could be decapped by enzymes similar to NUDT3, 12, or 15 which all yield RNAs with 5’-diphosphate ends [19]. Such RNAs are direct substrates for the cCE [16]. Finally, the RNMT-RAMAC heterodimer, which is directly bound to cCE, methylates the N^7^ position of capped RNA to yield an m^7^G cap that is indistinguishable from a nuclear-added cap [3].

Previous studies show that cCE maintains cap homeostasis, the cyclic decapping and recapping of target mRNAs [10, 20, 21]. Cap homeostasis is believed to regulate the stability of the target mRNAs and thus act as another post-transcriptional gene regulation hub [10]. Several live cell imaging studies have shown periodic translation events, and such behaviour is consistent with the cyclical decapping and recapping of mRNAs [22–24]. Further, cCE-targeted mRNAs have similar length poly(A) tails in both their capped and uncapped forms suggesting that recapped transcripts can be productively translated [25]. Despite these findings, blocking cytoplasmic capping for 24 hours had little discernible impact on steady state proteome complexity in cycling U2OS cells [26].

To date, cytoplasmic capping has only been studied in the context of mRNAs, and its activities on other pol II synthesized transcripts have not been explored. In this study, we examine if long non-coding RNAs (lncRNAs) are substrates for the cytoplasmic capping machinery. Like mRNAs, lncRNAs are transcribed by pol II and can be processed in multiple ways in the cytoplasm [27]. Transcriptome-wide studies identified different types of lncRNAs varying in size from 200 bp to 100 kb [28]. Similar to other pol II-transcribed RNAs, lncRNAs feature both 5’-caps and poly(A) tails [29]. Generally, lncRNAs contain both longer introns and fewer exons than mRNAs, but ~98% of lncRNAs are spliced [27]. Often lncRNAs fold into complex secondary structures which define their specificity toward their targets [30]. Many lncRNAs have tissue-specific expression patterns, but they are generally expressed at lower levels than mRNAs [28, 31].

LncRNAs have various functions such as regulating alternative splicing, nuclear trafficking, chromosomal architecture and chromatin remodelling, mRNA stability, and mRNA translation [32–36]. As shown by Cyrano, some lncRNAs are key nodes that regulate mRNA levels by interfering with specific microRNAs [37]. Another example, lncRNA ST3Gal6-AS1 binds histone methyltransferase MLL1, and thus functions as a guide by recruiting specific chromatin-modifying complexes to the promoter region of ST3Gal6 resulting in its transcription [38]. Further, lncRNAs such as MALAT1 act as scaffolds by providing surfaces to form ribonucleoprotein complexes [39]. Thus, depending on the biological cues, lncRNAs can function as positive or negative regulators of gene expression [40–43]. LncRNAs can also act as architectural RNA, or arcRNA [44]. For example, NEAT1, IGS, Hsromega, MALAT1, and SatIII initiate the formation of various membrane-less cellular bodies like stress bodies, nuclear speckles, paraspeckles, splicing speckles, and stress granules [44–50]. In addition, lncRNAs may participate in various stress response pathways by promoting liquid–liquid phase separation (LLPS) that result in alterations in the expression of RNA binding proteins [45]. For example, lncRNA NEAT1_2 expression is upregulated during various stresses and it promotes cell survival under such adverse conditions by sequestration of SFPQ protein in paraspeckles reducing SFPQ-mediated transcription of target genes [46].

LncRNA turnover is thought to mirror mRNA turnover by undergoing decapping, de-adenylation, and eventual exonucleolytic degradation [51]. Despite this assumption, the regulators and mediators of lncRNA turnover have not been studied methodically. In this study, we show that, like certain mRNAs, some lncRNAs are stable in an uncapped form. Further, our data show that several stable uncapped forms of lncRNAs accumulate in cells where cytoplasmic capping is inhibited, suggesting a subset of lncRNAs may be targeted by cytoplasmic capping.

## Materials and Methods

### Cell culture, plasmid constructs and transfection

The human osteosarcoma cell line (U2OS) was procured from ATCC. U2OS cells were grown in DMEM (Dulbecco’s Modified Eagle Medium, Himedia) supplemented with 10% Fetal Bovine Serum (Himedia) along with 1% penicillin-streptomycin (Invitrogen) in a humidified incubator in the presence of 5% CO2 at 37°C. pcDNA3-bio-myc-mCE was obtained from Addgene (#82475). The pcDNA4TO-bio-myc-mCE construct was prepared by sub-cloning the bio-myc-mCE sequence into an empty pcDNA4TO vector using KpnI and ApaI restriction sites. pcDNA4TO-bio-myc-K294A was prepared from pcDNA4TO-bio-myc-mCE by using In-Fusion Site Directed Mutagenesis kit (Takara Bio). The HA-NCK1-M3 construct was a generous gift from Dr. Louise Larose [52], and was prepared as described in Chen et al, 2000 [53]. Cells were transfected with either empty pcDNA4TO, pcDNA4TO-bio-myc-K294A, pRK5, or pRK5HA-NCK1-M3 using Jetprime transfection reagent (Polyplus) according to the manufacturer’s guidelines. Cells were harvested after 48 hours post-transfection.

### Cell fractionation

Harvested U2OS cell pellets were resuspended in cytoplasmic lysis buffer (20mM Tris-Cl of pH 7.5, 10mM NaCl, 10mM MgCl2, 10mM KCl, 0.2% NP40, 1mM PMSF, RNAseOUT and 1X protease inhibitor cocktail) and incubated on ice for 10 min with intermittent gentle agitation. Nuclei were removed by centrifugation at 1,000 xg for 10 minutes at 4°C. The nuclear pellet was resuspended in nuclear lysis buffer (150mM NaCl, 1% Triton X-100, 0.5% Sodium Deoxycholate, 1% SDS, and 25mM Tris-Cl of pH 7.5) for 40 minutes with occasional agitation. The nuclear extract was prepared by centrifugation at 10,000 xg for 10 minutes at 4°C.

### Western blotting

Equal amounts of nuclear and cytoplasmic protein fractions, as measured by BCA assay (Pierce), were loaded into 10% Mini-PROTEAN TGX gels (Biorad) and transferred onto Immobilon-FL PVDF (EMD Millipore) membranes. Membranes were blocked in 3% Bovine Serum Albumin in phosphate-buffered saline (PBS) for 30 minutes at room temperature. The blots were probed by adding primary antibodies, mouse anti p84 (Abcam #ab487), mouse anti-Lamin (DHSB #MANLAC1(4A7)), mouse anti-GAPDH (Novus #2D4A7), mouse anti-Myc (SCBT #32293), and rabbit anti-HA (Cell Signalling Technology, CST#3724)) at 1:1000 dilutions into the blocking solution overnight at 4°C. Membranes were rinsed three times in PBS +0.05% Tween-20 and probed with secondary antibodies, donkey anti-Mouse 800 (Invitrogen #SA535521) and donkey anti-Rabbit 680 (Invitrogen #A143) at 1:10000 dilutions in PBS +0.05% Tween-20. Results were visualized with an Odyssey CLx Imaging System (LiCor Inc.).

### Nuclear and cytoplasmic RNA extraction and poly(A) selection

Nuclear and cytoplasmic RNA were extracted from the respective fractions using Trizol reagent (Invitrogen) following the manufacturer’s protocol. RNAs were treated with DNase I (Invitrogen), as directed by the manufacturer and recovered using phenol: chloroform: isoamyl alcohol (SRL) extraction [25]. 2 μg of RNA from each fraction were poly(A) selected using Dynabeads mRNA DIRECT Kit (Invitrogen) as per manufacturer’s instructions.

### cDNA synthesis and PCR

Poly(A) selected RNAs were primed with oligo dT and reverse transcribed using a Super Script III First Strand cDNA Synthesis kit (Invitrogen). Primers were designed against selected lncRNAs from the 5’-ends for most transcripts except for RP11-574K11.28 and GAS5-838 where primers were designed against the 3’-ends. PCR was performed using Dream Taq DNA polymerase (Thermo Scientific). All oligonucleotides used in this study are listed in Table-1. The resulting PCR products were analyzed using ethidium bromide-stained 1.2% agarose gels, visualized with a Gel Doc (Biorad), and PCR amplicons were confirmed by Sanger sequencing.

**Table 1:**
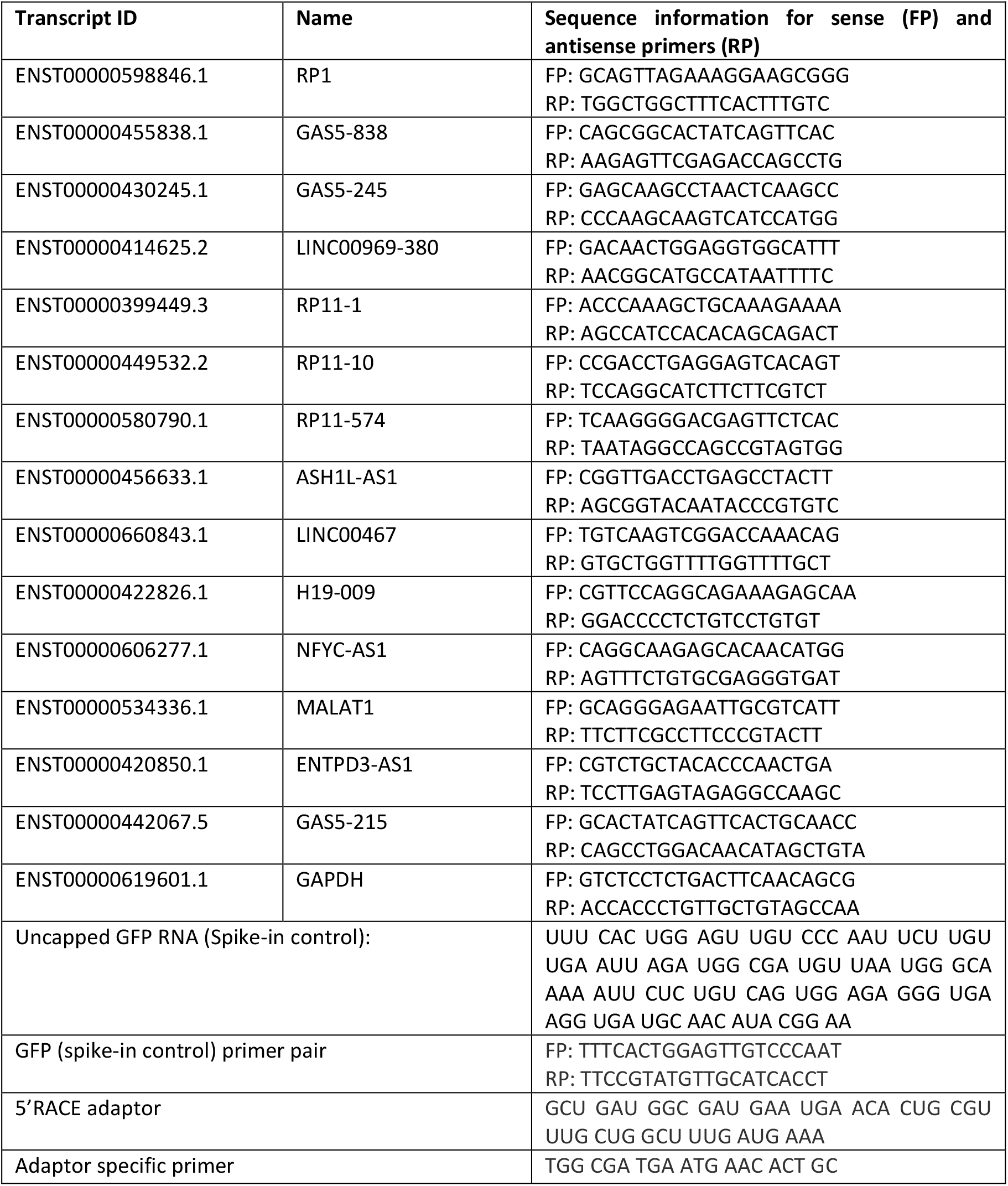
Sequences of primers and oligonucleotides used for PCR and 5’RACE assay.

### 5’-Rapid Amplification of cDNA Ends (5’-RACE)

5’-RACE was performed according to the previously published study as described in Mukherjee et al., 2012 [10]. Briefly, 36ng of heat-denatured poly(A)+ RNA was incubated at 37°C for 2 hours with 50 pmol/μl of RNA RACE adaptor and 10 units of T4 RNA ligase 1 followed by phenol-chloroform extraction and ethanol precipitation. The recovered RNA (~30 ng) was primed with random hexamers and reverse transcribed using Superscript III (Invitrogen) and analysed by semi-quantitative RT-PCR using an adaptor-specific forward primer and target gene-specific reverse primer. PCR products were separated using 1.2% agarose gels and visualized using the Biorad Gel-doc system. Certain PCR products were purified from the gel using NucleoSpin Gel and PCR Clean-up Mini kit (Macherey-Nagel) and cloned individually in a TA-cloning vector called pGEM®-T Easy Vector (Promega) according to the manufacturer’s instructions. Individual colonies were screened, and cloned sequence inserts were sent for Sanger sequencing to AgriGenome Labs Pvt Ltd. As spike in control, 100 bp GFP mRNA was synthesized with 5’-monophosphate ends (IDT). Sequence of this oligo is given in Table-1.

### CAGE analysis and CAGE databases

Cap analysis of gene expression (CAGE) queries were performed as described in [54], but modified to standardize each target’s genomic position to Hg19. Briefly, the annotated transcription start sites and cloned 5’-RACE ends of lncRNAs as shown in Table-3, were mined from the following predefined track All FANTOM5 CAGE libraries (n = 1897 pooled) (the track for H19 is loaded here, https://fantom.gsc.riken.jp/zenbu/gLyphs/#config=4liuCqdCRosxeSpjwKhcB;loc=hg19::chr11:2015741..2019730) during the week of September 26, 2022 using the ZENBU GENOME browser [55]. The genomic coordinates (aligned to Hg38) for the cloned 5’-RACE sequences were identified using the NCBI nucleotide BLAST function (https://blast.ncbi.nlm.nih.gov/Blast.cgi) searching the human (9606) NCBI taxonomy database. Hg38 build genetic coordinates were then converted to Hg19 build positions via the Lift Genome Annotation web tool (https://genome.ucsc.edu/cgi-bin/hgLiftOver) for visualization and subsequent CAGE analysis via the FANTOM5 database. Only CAGE tags mapping to the lncRNA designated strand (e.g., sense or antisense) directionality were retained for CAGE analysis.

**Table 2.** Selected lncRNAs for this study with additional identification terms for confirmation of target transcripts.

| Study Transcript Name | Alternative Transcript Name | Ensembl Transcript ID | LNCipedia Transcript ID | RefSeq Transcript ID | Length (bases) | Type |
| --- | --- | --- | --- | --- | --- | --- |
| RP1-283E3.8 | lnc-CDK11A-2 | ENST00000598846.1 | LNC-CDK11A-2:2 | Not annotated | 5079 | Intronic |
| GAS5-838 | GAS5-009 | ENST00000455838.1 | GAS5:10 | NR_002578<br>(partial overlap) | 632 | Bidirectional |
| GAS5-245 | GAS5-027 | ENST00000430245.1 | GAS5:1 | NR_002578<br>(partial overlap) | 723 | Antisense |
| LINC00969-380 | Lnc-MUC20-67 | ENST00000457233.1 | lnc-MUC20-67:1 | NR_003265<br>(partial overlap) | 1279 | Intergenic |
| RP11-464F9.11 | RP11-464F9.1-001 | ENST00000399449.3 | lnc-AGAP5-1:9 | Not annotated | 3810 | Intronic |
| RP11-464F9.12 | RP11-464F9.1-201 | ENST00000449532.2 | lnc-AGAP5-1:1 | NR_026592<br>(majority overlap) | 1131 | Intergenic |
| RP11-574K11.28 | RP11-464F9.1-003 | ENST00000580790.1 | lnc-AGAP5-1:2 | NR_026592<br>(majority overlap) | 1913 | Intergenic |
| LINC00467 | NR_026761.1 | ENST00000660843.1 | LINC00467:1 | NR_026761 | 758 | Intergenic |
| ASH1L-AS1 | RP11-29H23.1-002 | ENST00000456633.1 | ASH1L-AS1:3 | NR_027023<br>(partial overlap) | 911 | Antisense |
| H19-009 | H19 | ENST00000422826.1 | H19:11 | NR_002196<br>(majority overlap) | 1796 | Intergenic |
| NFYC-AS1 | NR_024567.1 | Not annotated | NFYC-AS1:1 | NR_024567 | 3182 | Antisense |
| MALAT1 | MALAT1-001 | ENST00000534336.1 | MALAT1:5 | NR_002819 | 8708 | Intergenic |
| ENTPD3-AS1 | ENTPD3-AS1-003 | ENST00000420850.1 | ENTPD3-AS1:1 | NR_040100<br>(partial overlap) | 791 | Intergenic |
| GAS5-215 | GAS5-018 | ENST00000442067.1 | GAS5:33 | NR_002578<br>(partial overlap) | 1114 | Intergenic |

**Table 3:**
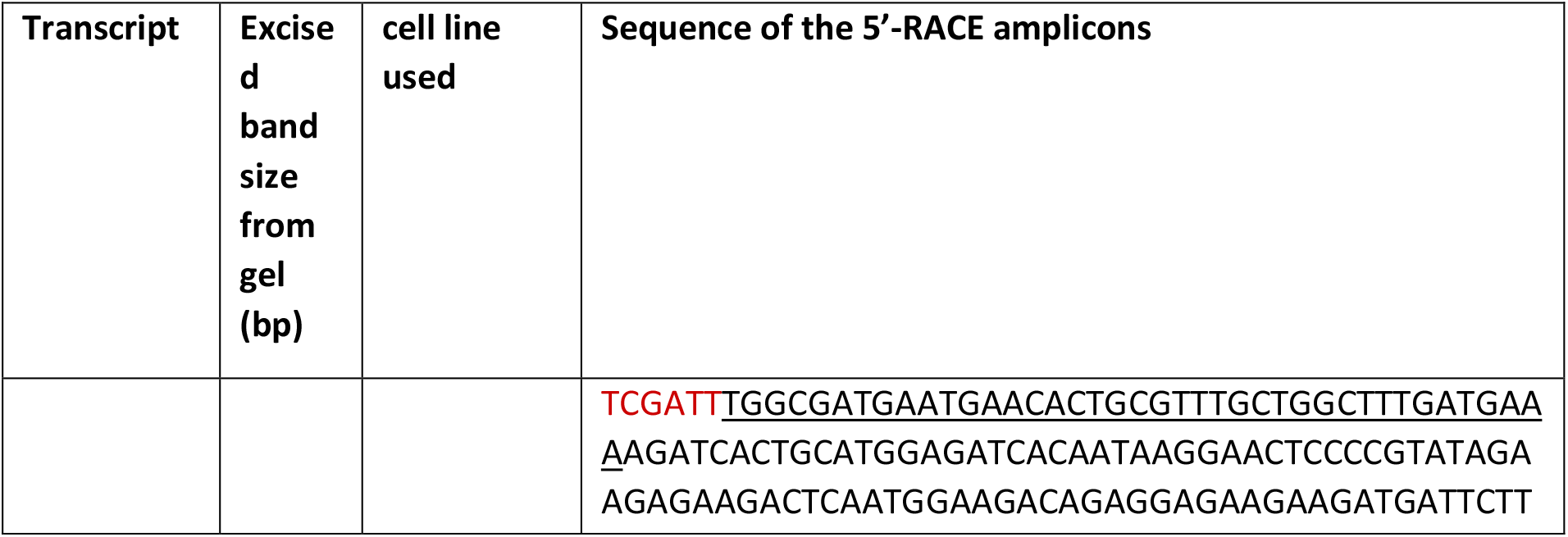
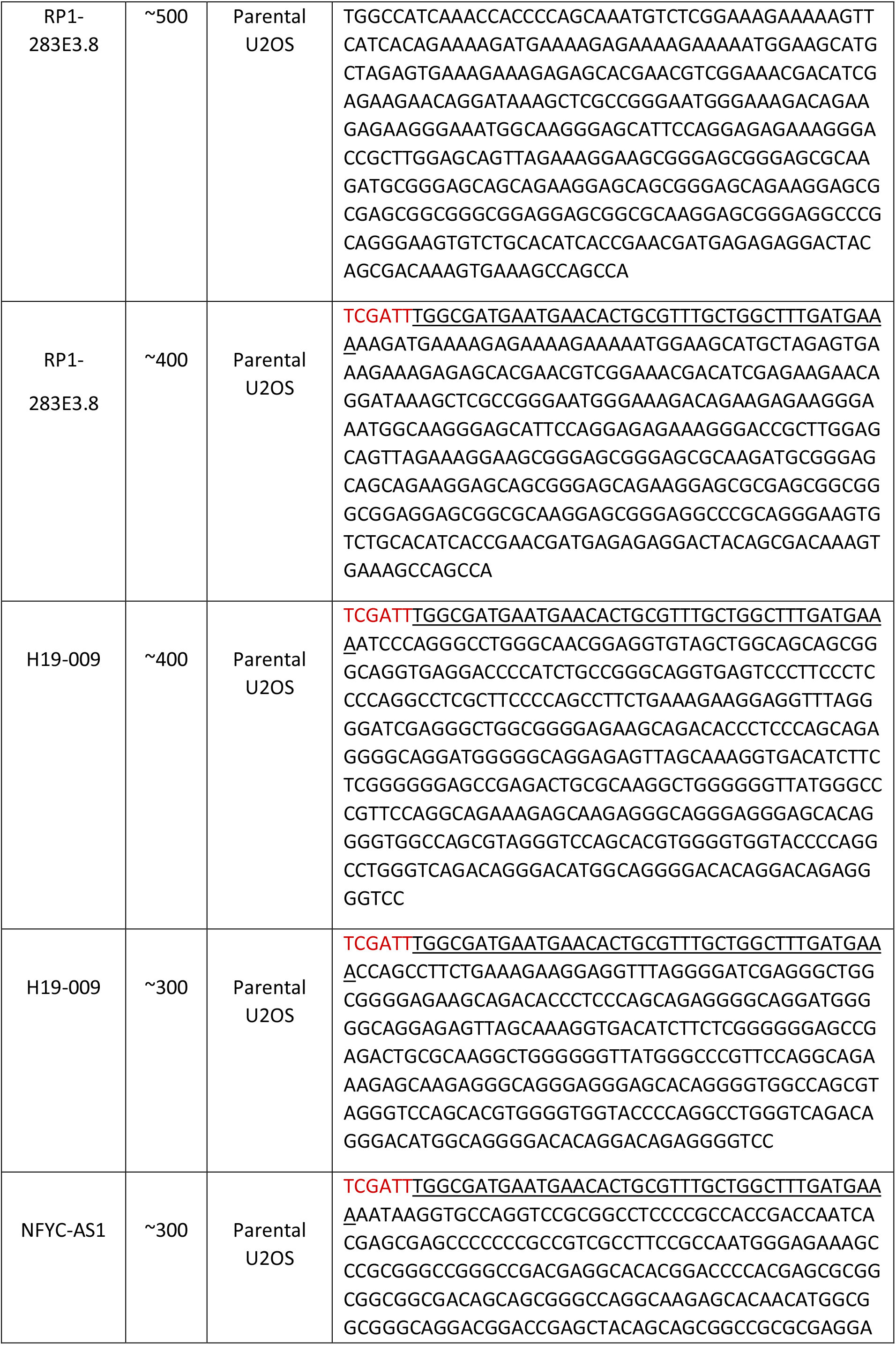
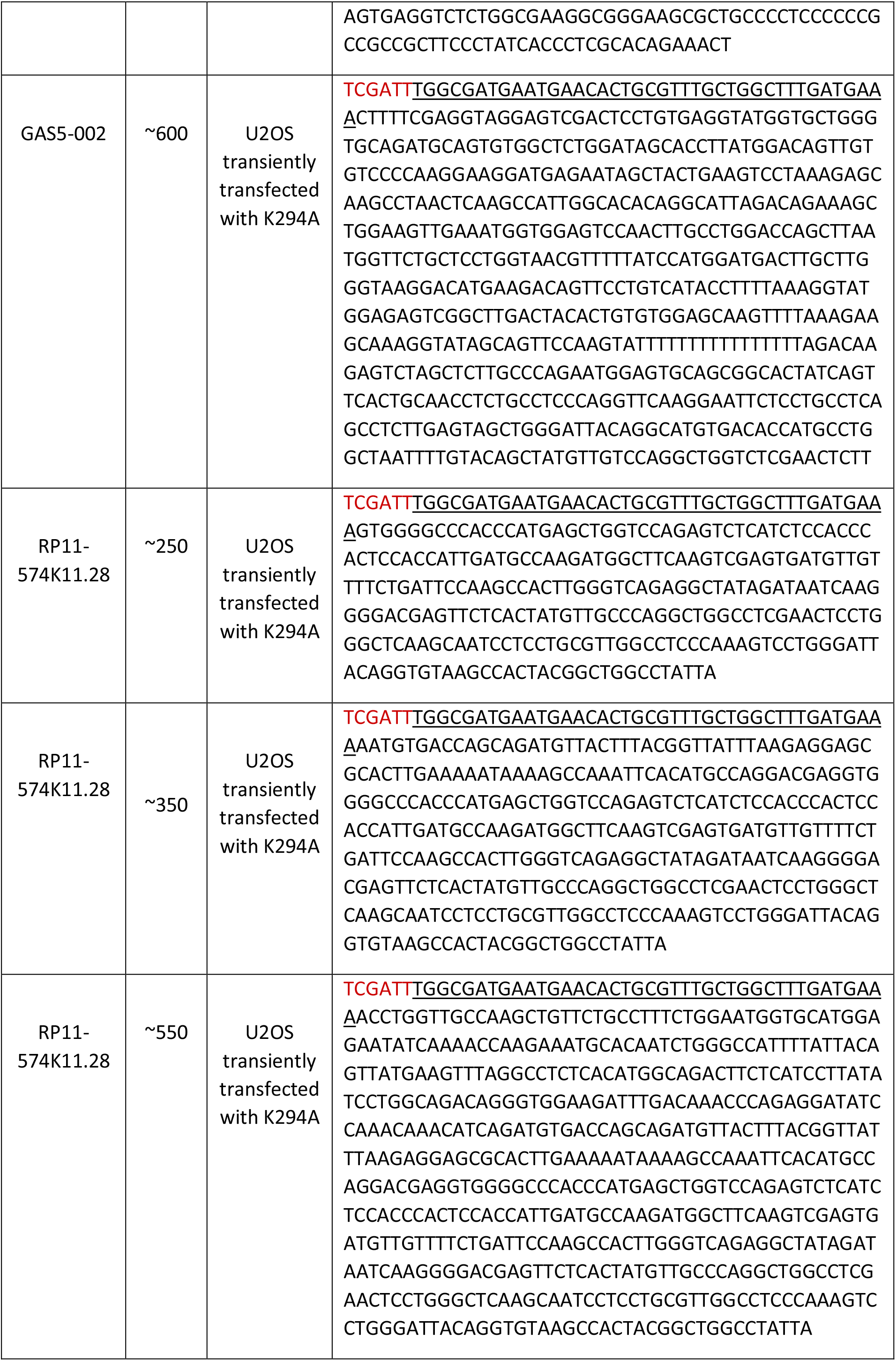
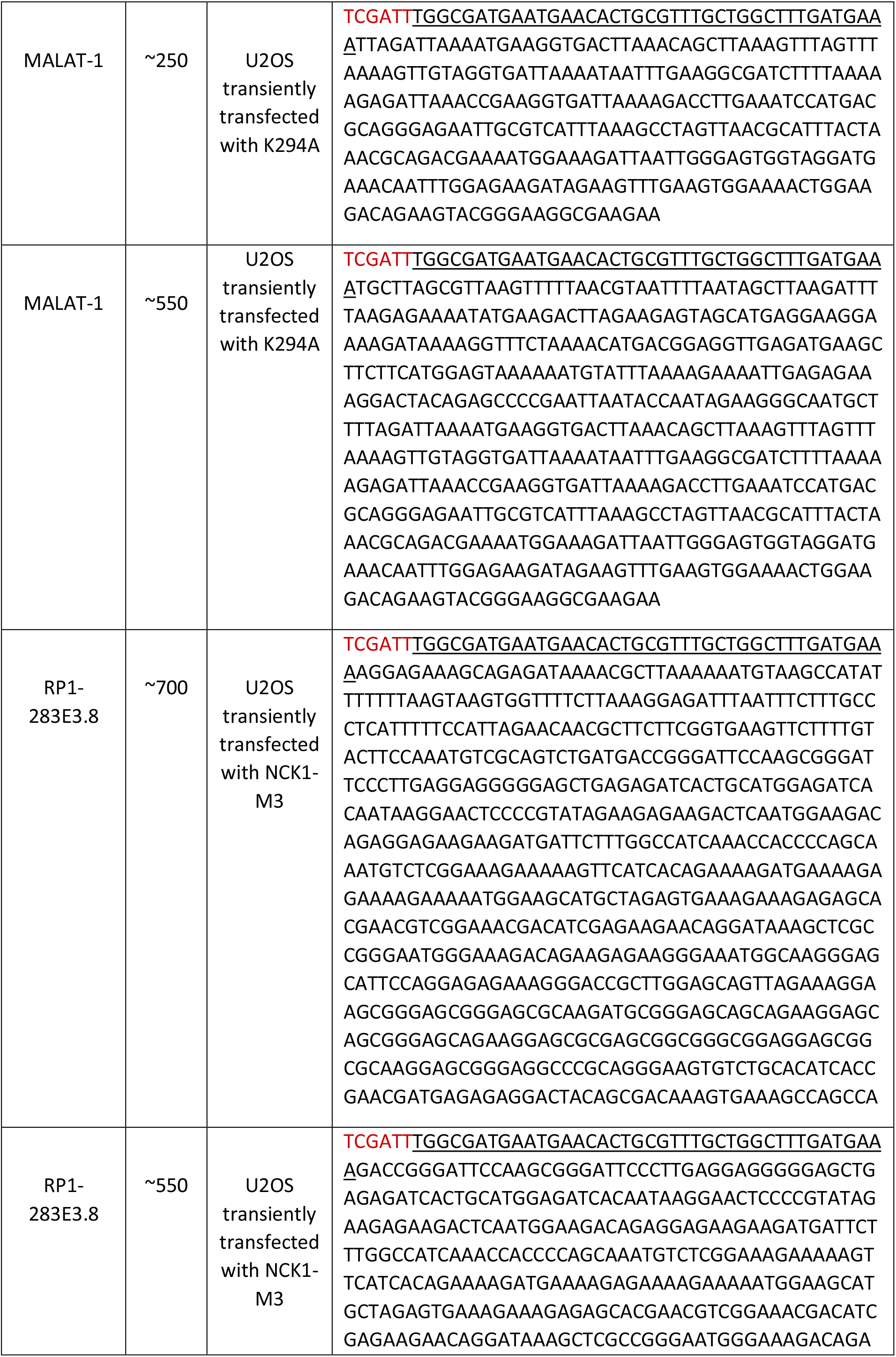
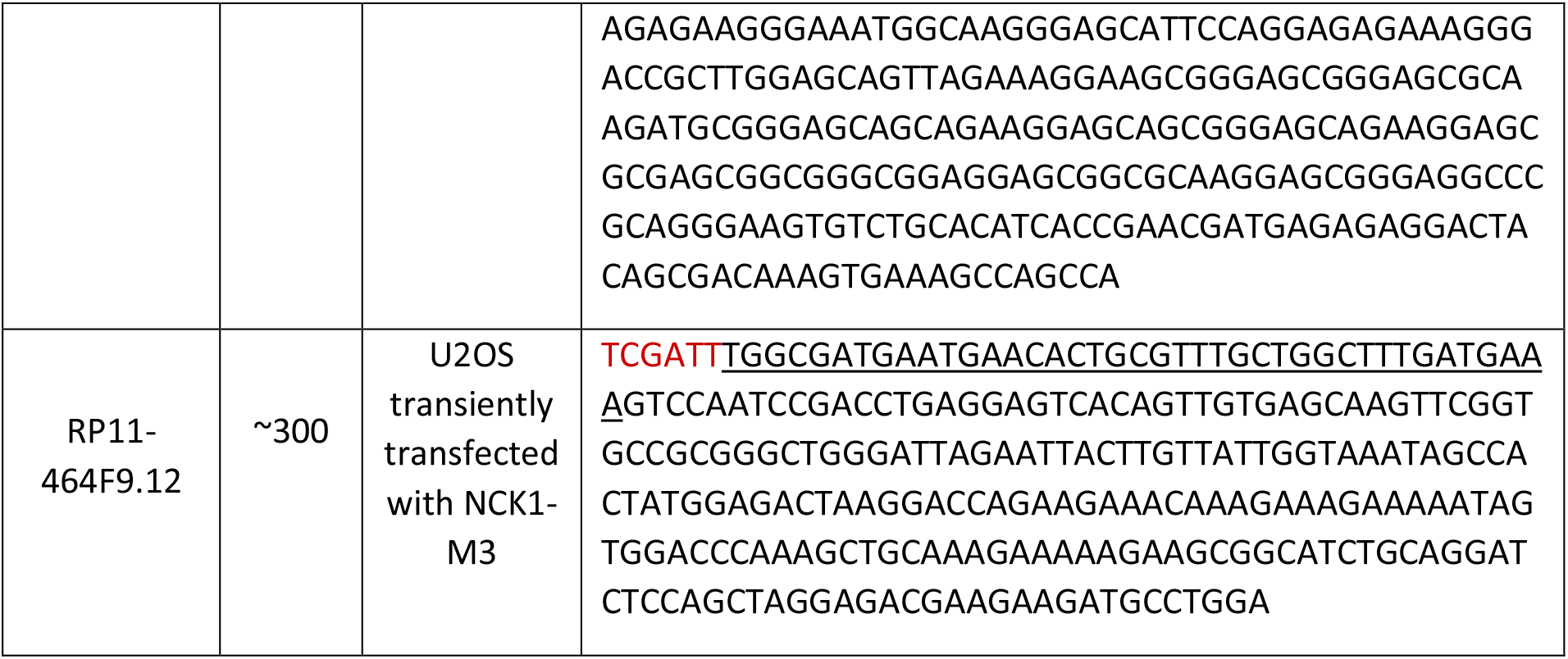
Sequences of the 5’-RACE amplicons of selected lncRNAs. Amplicons from parental U2OS cells and transfected cells transiently transfected with pcDNA4TO-Bio-Myc-K294A or pRK5-HANck1 M3. Here, the red colour signifies the part of the TA-cloning vector, underlined sequence denotes the RNA adaptor, and the rest indicates the candidate transcript with uncapped ends.

## Results and Discussion

### Nuclear and cytoplasmic localization of selected lncRNAs

To identify lncRNAs that might be regulated by cCE, we limited our *in silico* search for lncRNAs expressed in sarcomas since the cytoplasmic capping complex and its target mRNAs have been best characterized in U2OS osteosarcoma cells. CAGE uses the positions of m^7^G as a proxy for identifying transcription start sites (TSSs) [56]. Earlier studies with extensively validated mRNA targets of the cCE demonstrated that uncapped ends mapped exactly to, or in the vicinity of, downstream (non-TSS-associated) CAGE tags [54, 57]. We reasoned that lncRNAs could also be targets for the cCE complex. To test this hypothesis, we selected fourteen lncRNAs with non-TSS-associated as visualized by the UCSC genome browser (https://genome.ucsc.edu/cgi-bin/hgTracks) CAGE tags. The selected lncRNAs vary in size and classification as shown in Table-2. U2OS cells were fractionated into nuclear and cytoplasmic fractions to examine the subcellular localization of candidate transcripts. Efficient fractionation was validated by western blots using antibodies against nuclear (p84 and Lamin A/C) and cytoplasmic (GAPDH and α-tubulin) proteins (Fig. 1A). Our data show that the cytoplasmic fractions were devoid of nuclear contamination (Fig. 1A), and that each transcript was present in both nuclear and cytoplasmic fractions (Figs 1B and 1D). Here, our data shows most lncRNAs, like ASH1L-AS1, RP1, RP11 or different transcripts of GAS5, have similar nuclear and cytoplasmic localization (Figs 1B and 1D) except LINC00467 which is mostly present in the cytoplasm (Fig. 1C). Notably, our data confirm a previous report [58] showing that MALAT1, which is known to be associated with nuclear paraspeckles [59] is also present in the cytosol (Fig. 1D).

**Figure 1:**
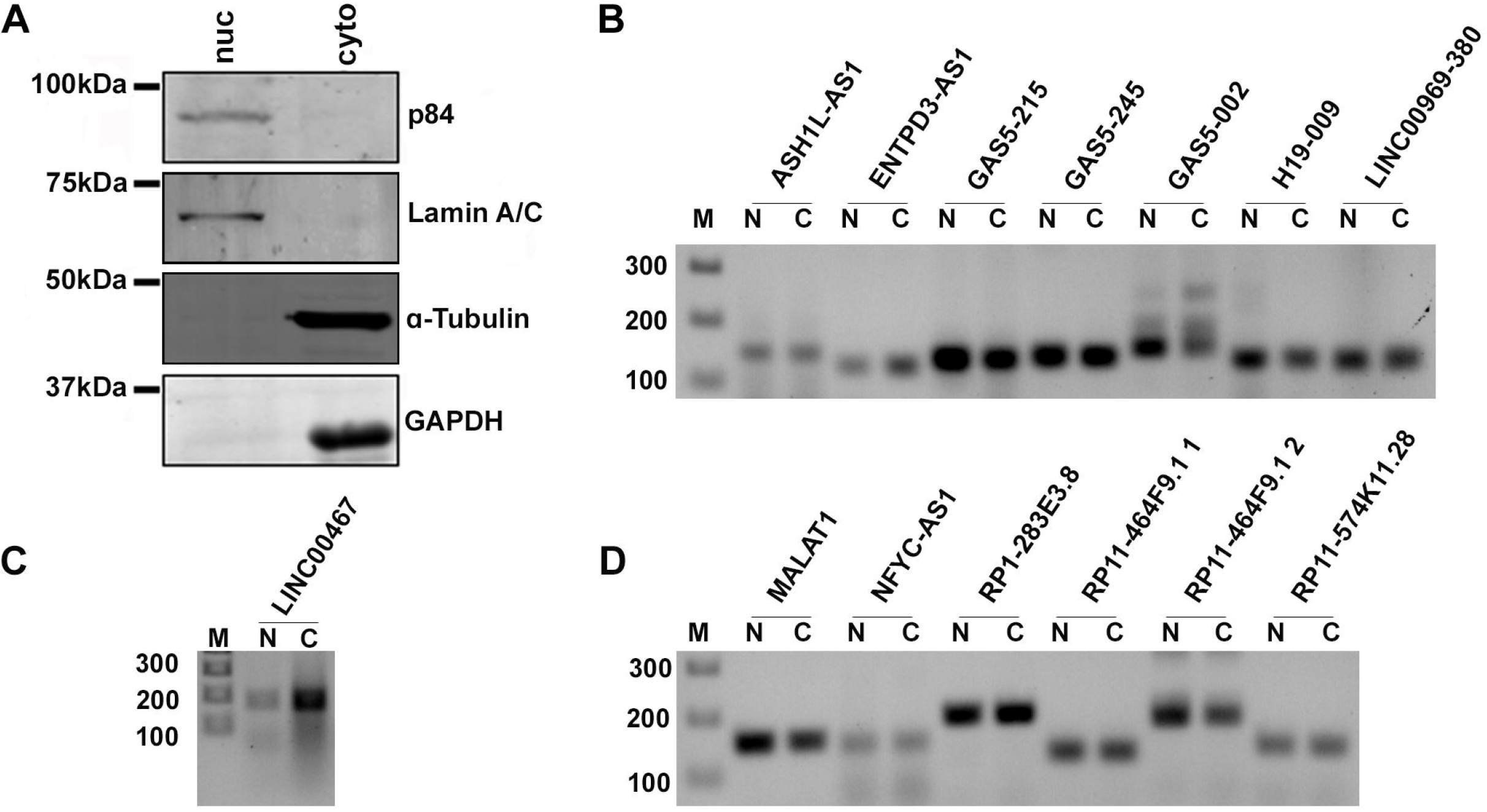
Nuclear and cytoplasmic localization of selected lncRNAs: (A) Cytoplasmic and nuclear protein extracts were prepared from U2OS cells. The quality of fractionation is demonstrated by Western blotting using antibodies against nuclear marker proteins p84 and Lamin A/C, and cytoplasmic marker proteins GAPDH and α-tubulin. The molecular weight of the proteins was indicated. (B-D) Nuclear and cytoplasmic RNA were extracted from the respective extracts. Detection of lncRNAs in nuclear and cytoplasmic fractions. N represents nuclear RNA and C represents cytoplasmic RNA, M: 100bp DNA marker.

### Detection of native uncapped forms of lncRNAs in U2OS cells

Most lncRNAs have poly(A) tails [28] which allowed us to recover them using oligo dT beads (Figs 2A-C). Only recovered cytoplasmic poly(A)-tailed RNAs were used as input for 5’-RACE assays. The ligation efficiency of the reaction was monitored using a synthetic RNA oligo encoding a portion of the GFP mRNA sequence as a spike in control as described earlier [10]. Our 5’-RACE assays detected both the spiked GFP RNA and VDAC3, an endogenous mRNA known to have uncapped transcripts (Fig. 2D). Our experiments detected multiple uncapped forms of candidate lncRNAs in native U2OS cells (Figs 2D-E). However, a few transcripts showed multiple uncapped forms but were recovered in lower amounts (Fig. 2E). Next, to validate if these amplicons were indeed generated from the 5’-RACE experiment, we excised the bands as shown in Fig. 2D-E from agarose gels and clone these products in TA vector for confirmation by sequencing. Sequencing data shows successful ligation of RNA adaptor with the uncapped ends of lncRNAs as shown in Figure 2F and Table-3.

**Figure 2:**
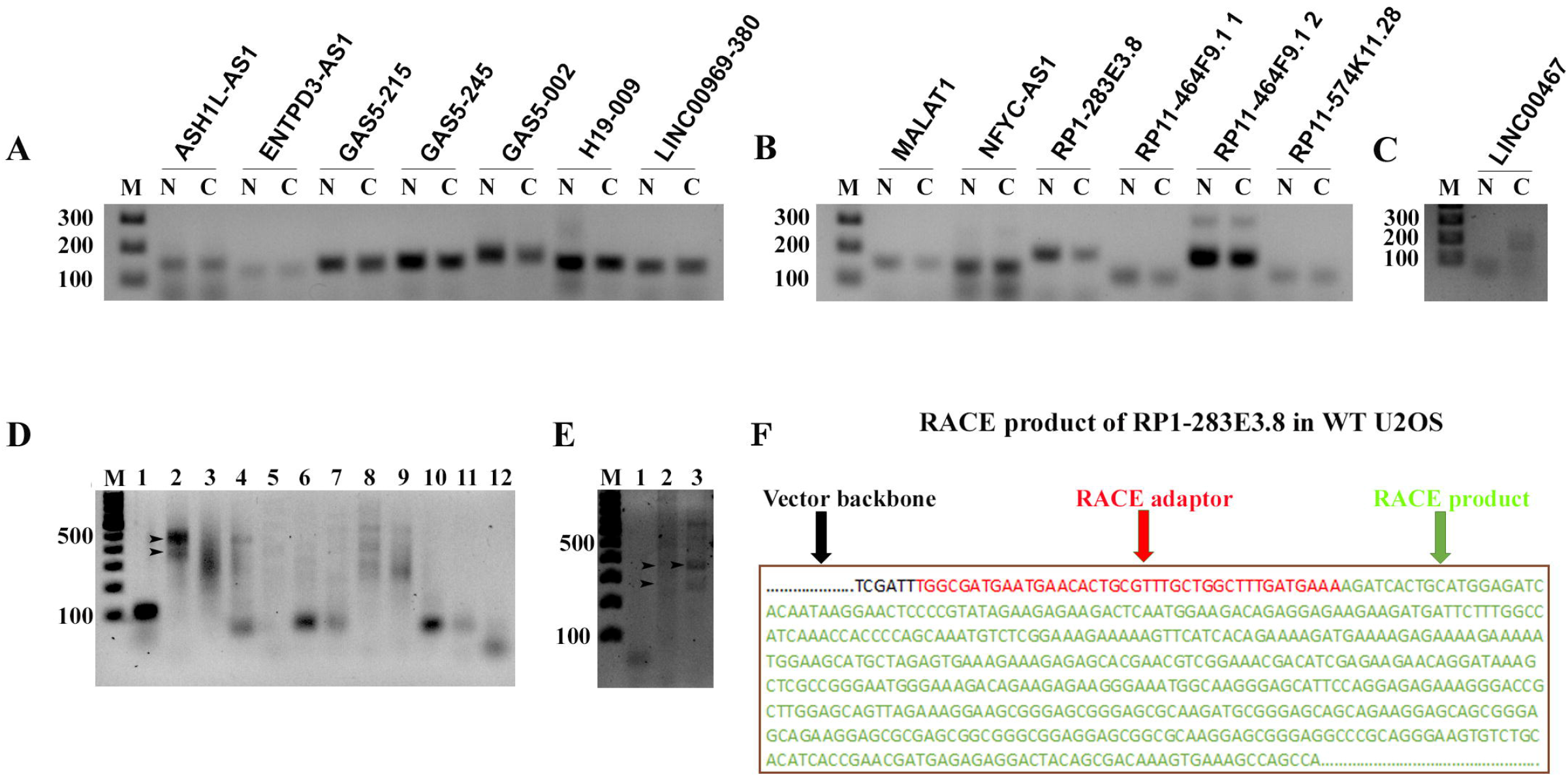
Detection of native uncapped forms of lncRNAs. (A-C) Detection of lncRNAs in poly (A) containing RNAs isolated from nuclear (N) and cytoplasmic (C) RNAs. (D-E) The presence of uncapped ends of candidate lncRNAs in parental U2OS cells was revealed by 5’RACE technique. The lanes are shown below, (D) lane 1: GFP spike in control, lane 2: RP1-283E3.8, lane 3: GAS5-002, lane 4: GAS5-245, lane 5: LINC00969-380, lane 6: RP11-464F9.1 1, lane 7: RP11-464F9.1 2, lane 8: RP11-574K11.28, lane 9: MALAT-1, lane 10: LINC00467, lane 11: ASH1-AS1, lane 12: VDAC3 and (E) lane 1: ENTPD3-AS1, lane 2: H19, lane 3: NFYC-AS1 (D). M represents 100 bp marker. Arrows in the figures indicate the amplicons that are excised from gel, cloned in TA vector and sent for Sanger sequencing. (E) Example sequence obtained from RACE product generated by RP1-283E3.8 from parental U2OS cells is shown. Vector sequence, adaptor sequence and uncapped ends with the RACE product are shown as indicated.

### Uncapped forms of lncRNAs accumulate in cytoplasmic capping inhibited cells

Earlier studies have shown mRNAs with uncapped ends can accumulate in cells where cytoplasmic capping was blocked [10, 54, 57]. The presence of uncapped lncRNAs led us to examine if these transcripts are targeted by cytoplasmic capping complex. We developed a cytoplasm-restricted, N terminally biotin and Myc tagged dominant negative mutant of cCE (K294A) [16], to study cytoplasmic capping independent of nuclear capping. U2OS cells were transiently transfected with empty vector control pcDNA4TO or pcDNA4TO-bio-myc-K294A (Fig. 3A) and the 5’-RACE experiments were repeated as before. Equal recovery of GFP spike-in control from control cells and K294A expressing cells indicated similar RNA ligation efficiencies in both sets of experiments (Fig. 3D, lanes 1-2). Interestingly, several transcripts like ENTPD3-AS1, H19-009, RP1-283E3.8 were more prevalent in cells expressing epitope-tagged K294A compared to control cells (Figs 3C-E). Our data also validate earlier results showing that uncapped VDAC3 mRNA is more detectable in cytoplasmic capping-inhibited cells (Fig. 3D, lanes 3-4) which was reported earlier as a target of cytoplasmic recapping [10, 54, 57].

**Figure 3:**
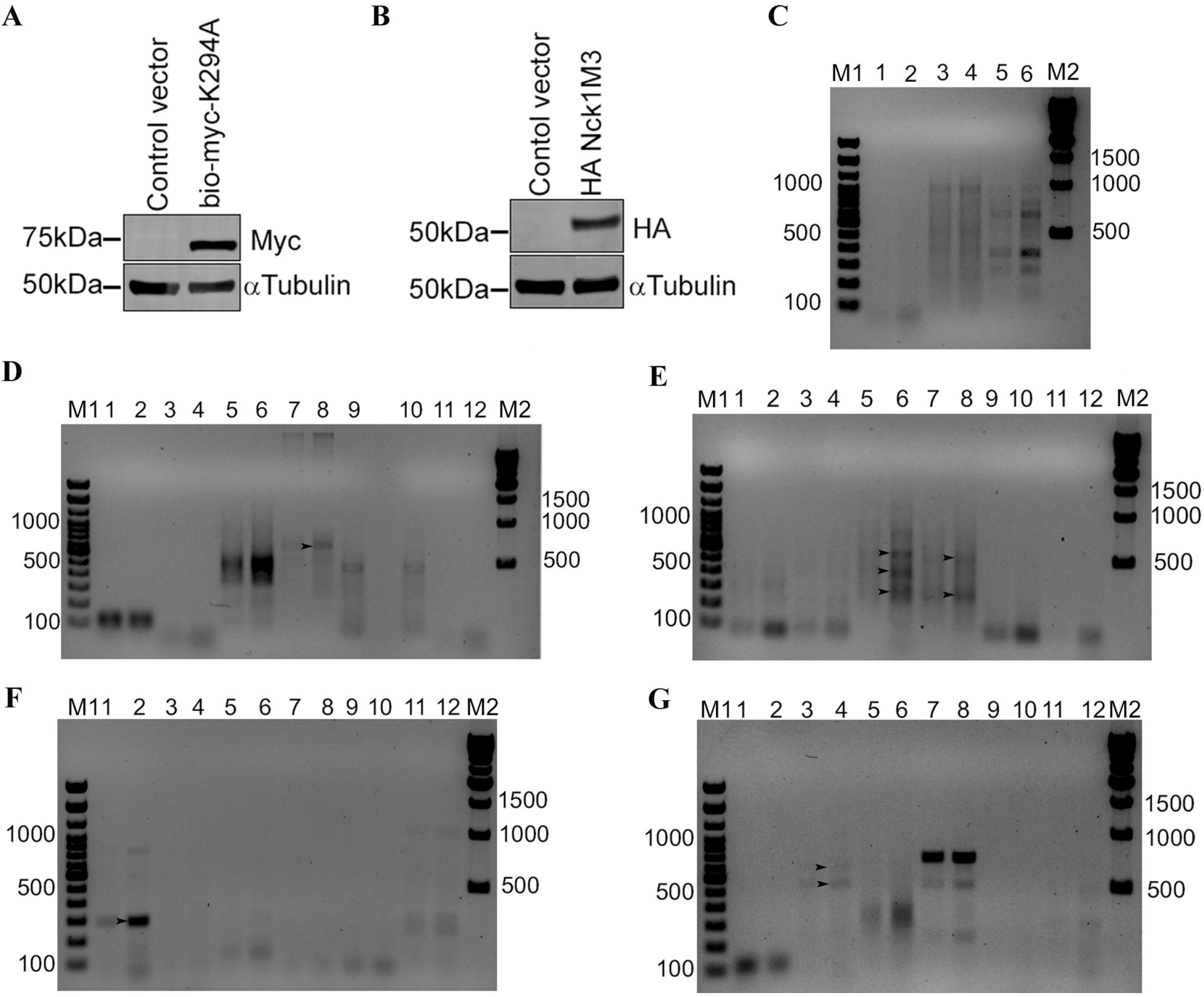
Accumulation of native uncapped lncRNAs in cytoplasmic capping inhibited cells. (A-B) The exogenous expression of bio-myc-K294A and HA- Nck1M3 are analysed by western blotting with anti-Myc and anti-HA antibodies on cytoplasmic protein extracted from transiently transfected cells with constructs encoding Bio-Myc-K294A and HA-Nck1 M3. Anti-alpha-Tubulin antibody is used to detect equal amount of protein loaded in each panel. The molecular weight of the proteins was indicated. (C-E): The presence of uncapped ends of candidate lncRNAs in U2OS cells either transfected with empty vector (odd lanes) or pcDNA4TO-Bio-Myc-K294A (even lanes). The lncRNAs are indicated as, C) lane 1,2: ENTPD3-AS1, lane 3,4: H19-009, lane 5,6: NFYC-AS1. (D) lane 1,2: GFP spike-in control, lane 3,4: VDAC3 mRNA, lane 5,6: RP1-283E3.8, lane 7,8: GAS5-002, lane 9,10: GAS5-245, lane 11,12: LINC00969-380. (E) lane 1,2: RP11-464F9.1 1, lane 3,4: RP11-464F9.1 2, lane 5,6: RP11-574K11.28, lane 7,8: MALAT-1, lane 9,10: LINC00467, lane 11,12: ASH1L-AS1. (F-G) The presence of uncapped ends of candidate lncRNAs in U2OS cells either transfected with empty vector (lanes 1,3,5,7,9,11) or pRK5-HANck1 M3 (lanes 2,4,6,8,10,12). The lncRNAs are indicated as, (F) lane 1,2: RP11-464F9.1 2, lane 3,4: RP11-574K11.28, lane 5,6: MALAT-1, lane 7,8: ASH1L-AS1, lane 9,10: H19-009, lane 11,12: NFYC-AS1. (G) lane 1,2: GFP spike-in control, lane 3,4: RP1-283E3.8, lane 5,6: GAS5-002, lane 7,8: GAS5-245, lane 9,10: LINC00969-380, lane 11,12: RP11-464F9.1 1. M1 and M2 represent 100 bp and 500 bp marker respectively. Arrows in the figures indicate the amplicons that are excised from gel, cloned in TA vector and sent for Sanger sequencing.

We leveraged the structure of the cytoplasmic capping complex to block cytoplasmic capping in an independent manner as a way to confirm that the accumulation of uncapped lncRNAs was caused by blocking cytoplasmic capping. As stated earlier, the Nck1 protein serves as the scaffold for the cytoplasmic capping complex and blocking the interaction of cCE with Nck1 also prevents cytoplasmic capping [18]. U2OS cells were transiently transfected with either empty vector or an inactive mutant of HA-Nck1 protein (HA-Nck1-M3) that cannot bind cCE (Fig. 3B) [18]. Cytoplasmic RNA was extracted from control and HA-Nck1-M3 cells overexpressing cells and 5’-RACE assays were then performed as before. Consistent with a role of the cytoplasmic capping complex in stabilizing these uncapped RNAs, increases in uncapped forms of several transcripts were evident in cells overexpressing Nck1-M3 compared control (Figs 3F-G). Select amplicons were sequenced and several different processed intermediates were revealed (Table-3). Together, these complimentary results link the cytoplasmic capping complex to the increased prevalence of uncapped forms of selected lncRNAs like H19-009, NFYC-AS1, RP1-283E3, GAS5-002, GAS5-245, LINC00969-380, RP11-464F9.11, RP11-464F9.12, RP11-574K11.28, MALAT-1, LINC00467, and ASH1L-AS1.

### Association of CAGE tags with putative cytoplasmic recapping sites

The transcripts of fourteen lncRNAs were targeted to determine the potential role of cytoplasmic capping in the post-transcriptional gene regulation in U2OS cells. The capture of select uncapped lncRNA transcripts by 5’-RACE was evaluated under three conditions in U2OS cells: the parental (empty expression vector) U2OS cells, and those transiently transfected with either K294A or Nck1-M3 for inhibition of the cCE. The location and number of CAGE tags within a narrow range (±60 bases) of the 5’-RACE-mapped end are shown in Supplemental Figures 1–2. Position 0 indicates the transcript’s 5’-RACE-mapped end and the number of CAGE tags observed at that particular genomic position are shown.

We feature three transcripts in Figure 4 to highlight the relationship between the uncapped state of these transcripts and the number of CAGE tags observed at the 5’-ends of the RACE-cloned sequences. For H19-009 Product 1 (Fig. 4A), 991 CAGE tags, which is ~2-fold greater than the surrounding region, map to the transcript’s RACE-detected 5’-end. Next, in cells transfected with the inactive K294A form of cCE, 5’-RACE recovered the uncapped 5’-end of the full-length GAS5-002 transcript. Further, we observed a tight grouping of CAGE tags mapping at the isolated RACE product for GAS5-002 (Fig. 4B). Importantly, although the GAS5-838 transcript (ENST00000455838.1) was targeted, the reverse primer captured and amplified GAS5-002 (ENST00000431268.1), which has overlapping genomic coordinates with the intended target. A similar cluster of CAGE tags is observed for the final transcript, RP11-464F9.12 from Nck1-M3 transfected cells. 6,066 CAGE tags map to the 5’-end of the recovered 5’-RACE product (Fig. 4C). Importantly, in both cases where cytoplasmic capping was blocked, our RACE experiment detected the native, uncapped, 5’-end of the targeted lncRNA. Those results suggest that the cytoplasmic capping complex maintains the cap on these lncRNAs through cap homeostasis [10].

**Figure 4:**
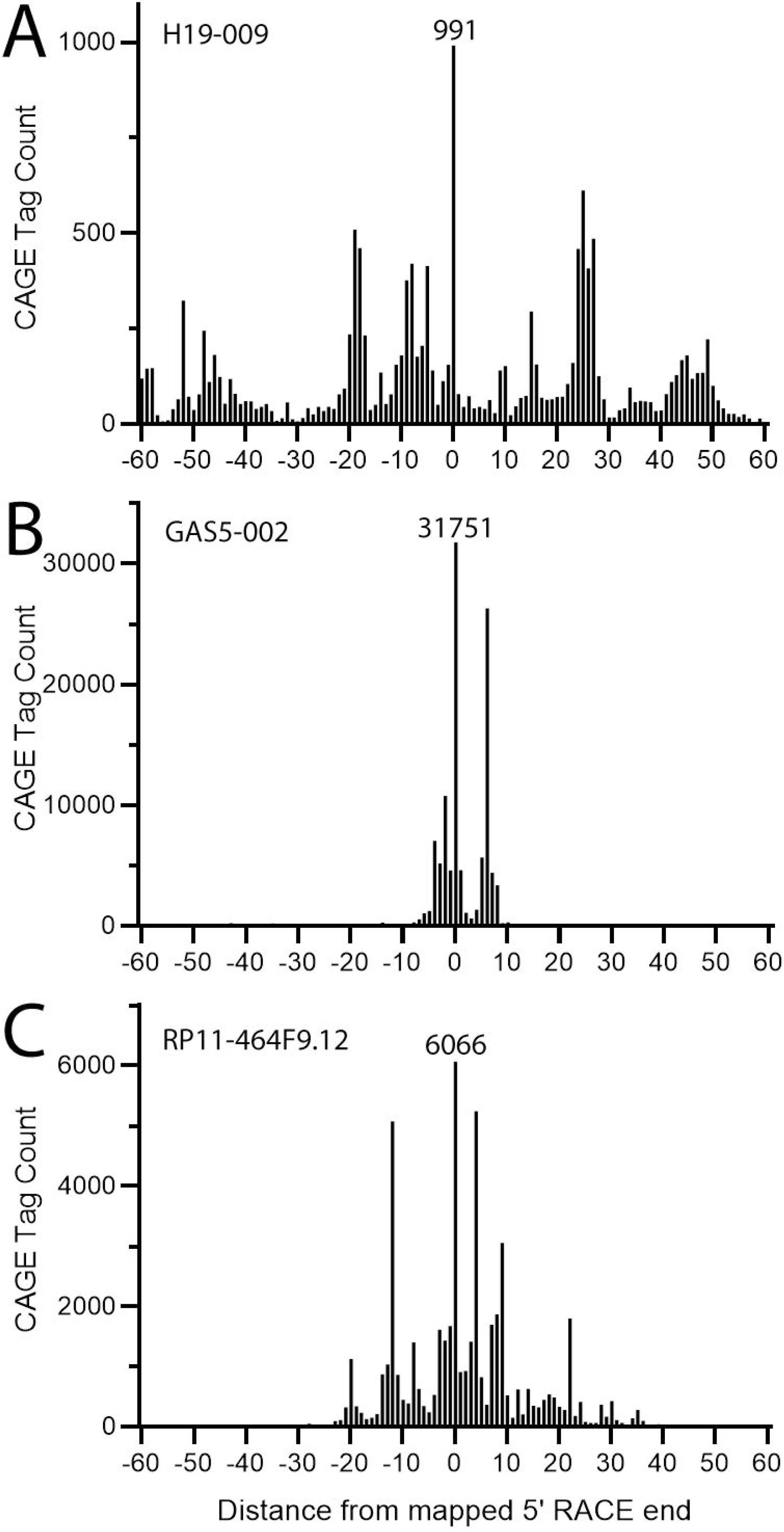
CAGE tags surrounding the 5’-end of transcripts captured by 5’-RACE under various transfection conditions. The 5’-end sequences of uncapped candidate transcripts were detected by 5’-RACE. Positions and associated CAGE tags are shown in ±60 base stretches of the 5’-end (position 0) of the 5’-RACE captured transcripts for (A) H19-009 Product 1 (parental U2OS cells), (B) GAS5-002 Product (K294A transiently transfected cells), and (C) RP11-464F9.12 Product (Nck1-M3 transiently transfected cells).

Notably, we saw instances where the targeted lncRNA transcript shared genomic coordinates with a protein-coding mRNA (RP1-283E3.8 (ENST00000598846.1) overlapping with CDK11A for example). Three of the RP1-283E3.8-targeting products captured by 5’-RACE (Figs S1A, S1B and S2G) map to the CDK11A mRNA sequence, not the targeted lncRNA. Capture of the mRNA was confirmed when the captured sequences were shown to include a CDK11A mRNA-specific splice junction. True RP1-283E3.8 lncRNAs possess an extension in exon 3 that is not present in CDK11A mRNA, but is observed for RP1-283E3.8, Product 1 in the Nck1-M3 transfected cells (Fig. S2F). However, as the captured sequences overlap with protein-coding mRNA sequences we cannot unequivocally assign the CAGE tags to either the lncRNA or mRNA.

Importantly, this narrow window of CAGE data only provides a limited understanding of the possible cap positions on a lncRNA. The number of CAGE tags in any given peak is best understood in the context of how a particular CAGE peak compares to the CAGE tags mapping across the full-length transcript (Fig. S3). When examined in this context, the highest CAGE peaks mapping to the narrow regions of H19 is less than 10% of the full-length transcript’s highest CAGE tag peak near the TSS (Fig S3A). The full-length CAGE analysis confirms that, in general, most CAGE tags are localized to the 5’-ends of the lncRNAs we studied (Figs S3B and S3C).

Taken together, our results show that a population of select lncRNAs are present in an uncapped state and that some of these could be targeted by the cytoplasmic capping machinery. Further, we detected multiple uncapped forms of lncRNAs and also precisely mapped the positions of their uncapped ends with single nucleotide resolution. The generation of one or multiple uncapped forms of lncRNAs in cytoplasmic capping inhibited cells by overexpressing K294A and Nck1-M3 suggested that these lncRNAs could be processed differently and could possibly be recapped at different sites. Future studies will be directed to reveal the role of cap homeostasis in regulating these lncRNAs. For example, analyses using CAGE, teloprime [60] or related techniques coupled with degradome sequencing [61] can validate the sequences obtained via RACE experiments and/or identify new candidate cytoplasmic capping sites. In addition, future studies should be directed to identify specific sequences that might protect these lncRNAs from cellular nucleases and/or target these lncRNAs to the cytoplasmic capping machinery.

## Acknowledgements

This work was supported by extramural grant CRG/2019/006427 from Department of Science and Technology, Government of India to Dr. Chandrama Mukherjee. The authors thank SERB-CRG for providing fellowship to Mr. Avik Mukherjee. The authors also thank UGC, Government of India for a UGC-SRF fellowship to Mr. Safirul Islam and Department of Biotechnology, India for a Ramlingaswami Re-entry fellowship to Dr. Chandrama Mukherjee. The authors thank Presidency University for necessary infrastructural support, Dr. Piyali Mukherjee, Presidency University for providing Lamin antibody, Dr. Shubhra Majumder, Presidency University for providing HA antibody and Dr. Abhik Saha, Presidency University for providing access to Licor imager. The Kiss Lab’s contributions are supported by a grant from the National Institutes of Health (R35GM137819, to DLK) and Dr. Rachel Kieser is supported by a postdoctoral supplement (R35GM137819-03S3).

## Author contributions

CM conceived, designed and supervised the experiments for AM and SI. AM and SI performed the experiments described in study. REK performed the CAGE analysis with guidance and supervision from DLK. AM, SI, REK, DLK, and CM all wrote different sections of the manuscript. All authors contributed to editing and revising the manuscript, and all authors have approved the final manuscript. The authors have no conflict of interests.

