## Supplementary Information for "Identification of long non-coding RNAs as substrates for cytoplasmic capping enzyme"

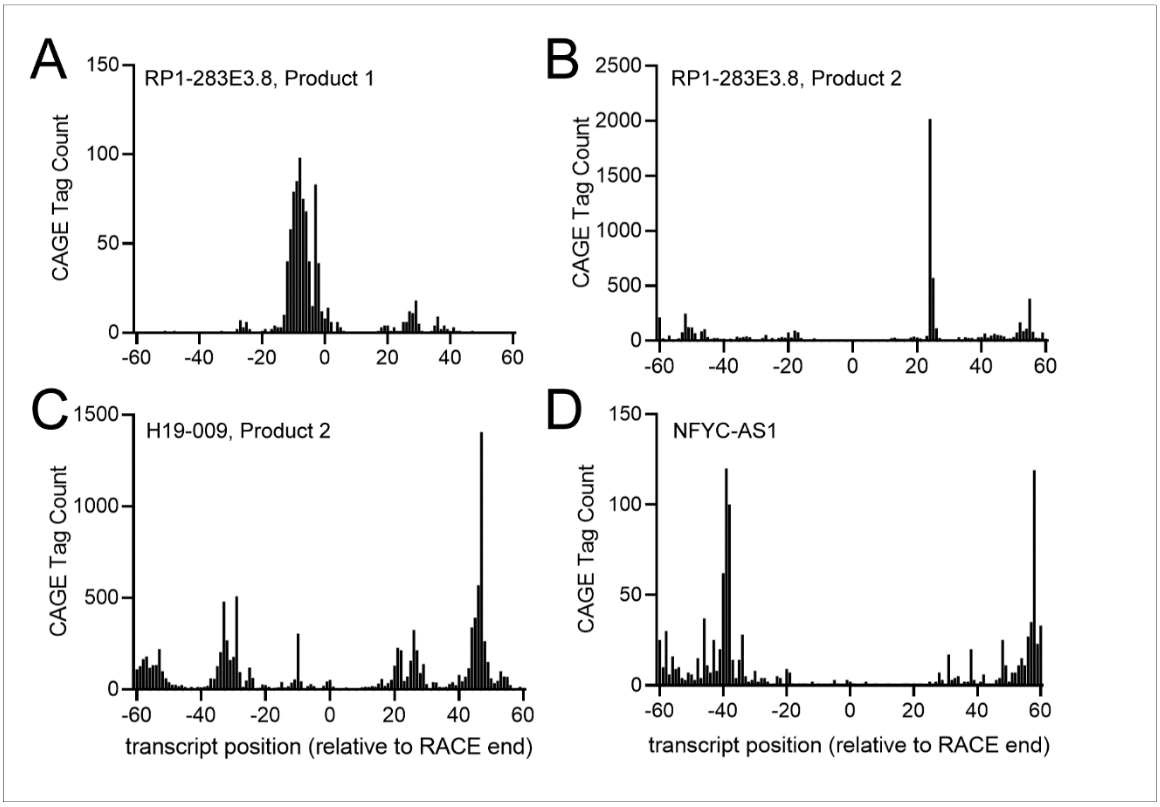


**Supplemental Figure 1: CAGE tags near the 5´-RACE ends from the parental U2OS cells.** Uncapped 5´-end sequences were detected by 5´-RACE. Positions and number of CAGE tags are shown in ±60 base stretches of the 5´-end (position 0) of the 5´-RACE captured transcripts for (A) RP1-283E3.8 Product 1, (B) RP1-283E3.8 Product 2, (C) H19-009 Product 2, and (D) NFYC-AS1.

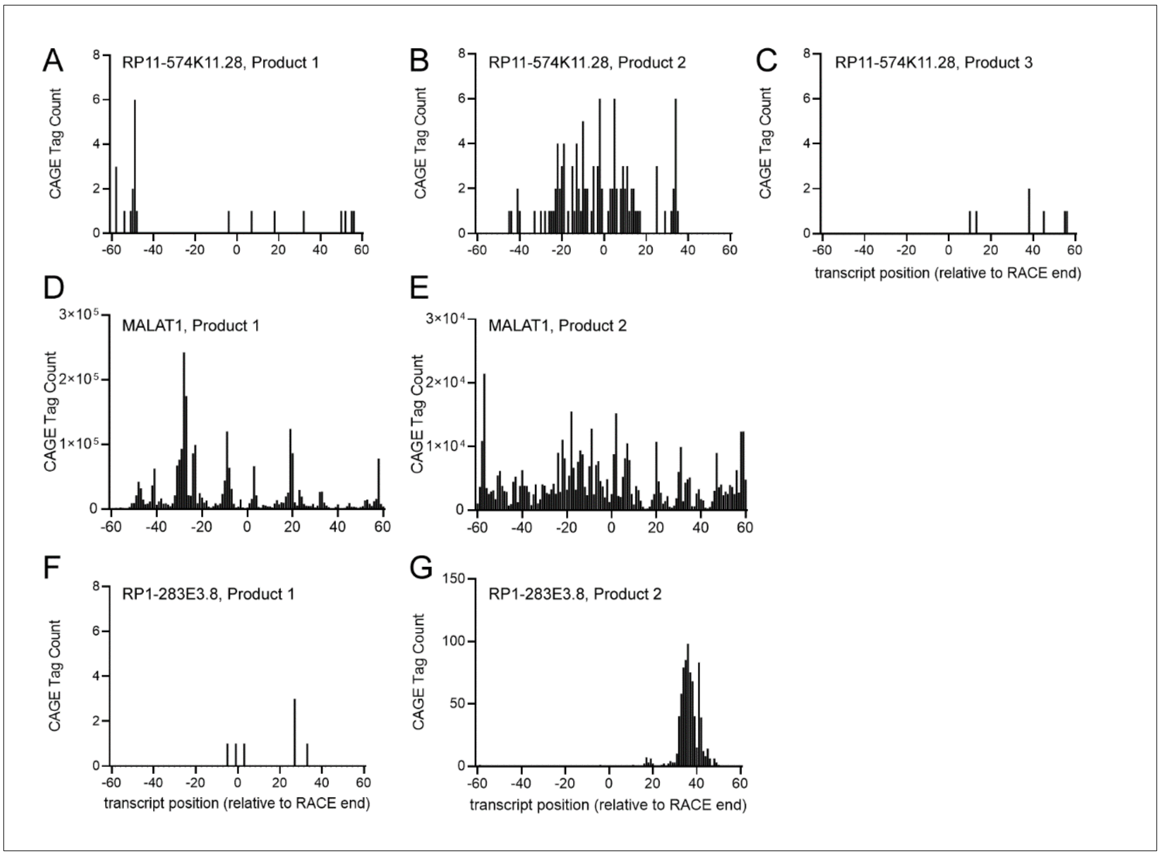


**Supplemental Figure 2: CAGE tags near the 5´-RACE ends from U2OS cells transiently transfected to block the cytoplasmic capping enzyme complex.** Uncapped 5´-end sequences were detected by 5´-RACE. Positions and number of CAGE tags are shown in ±60 base stretches of the 5´-end (position 0) of the 5´-RACE captured transcripts. The CAGE tags for transcripts from cells transiently transfected with pcDNA4TO-Bio-Myc-K294A are shown for (A) RP11-574K11.28 Product 1, (B) RP11-574K11.28 Product 2, (C) RP11-574K11.28 Product 3, (D) MALAT-1 Product 1, and (E) MALAT-1 Product 2. In addition, the CAGE tags for transcripts from cells transiently transfected with pRK5-HANck1 M3 are shown for (F) RP1-283E3.8 Product 1 and (G) RP1-283E3.8 Product 2.

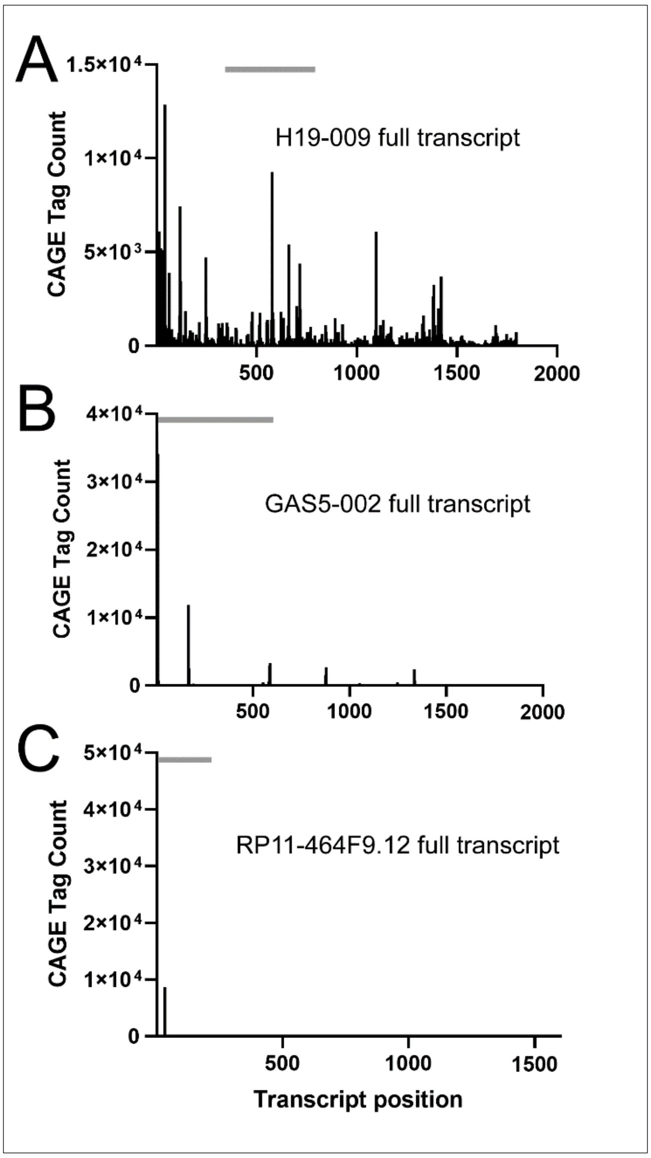


**Supplemental Figure 3: CAGE results for full-length lncRNA transcripts.** The positions along the transcript (identified in sequential order according in the 5´-to-3´ direction on the antisense strand) and the number of CAGE tags are shown across the length of the lncRNA transcript. Due to the considerable length of the selected lncRNA transcripts, the CAGE tags have been binned; therefore, the CAGE tags have been combined for a total tag count across a particular base pair range. The bar above the CAGE tags indicates the approximate positioning of the 5'-RACE captured cloned-sequences in Figure 4. The CAGE results for the full-length transcripts for (A) H19-009 (total length: 1,796 bps with 3 bp binning), (B) GAS5-002 (total length: 1,697 bps with 3 bp binning), and (C) RP11-464F9.12 (total length: 1,131 bp with 32 bp binning) are shown.
